# Phylodynamics on local sexual contact networks

**DOI:** 10.1101/082966

**Authors:** David A. Rasmussen, Roger Kouyos, Huldrych F. Günthard, Tanja Stadler

## Abstract

Phylodynamic models are widely used in infectious disease epidemiology to infer the dynamics and structure of pathogen populations. However, these models generally assume that individual hosts contact one another at random, ignoring the fact that many pathogens spread through highly structured contact networks. We present a new framework for phylodynamics on local contact networks based on pairwise epidemiological models that track the status of pairs of nodes in the network rather than just individuals. Shifting our focus from individuals to pairs leads naturally to coalescent models that describe how lineages move through networks and the rate at which lineages coalesce. These pairwise coalescent models not only consider how network structure directly shapes pathogen phylogenies, but also how the relationship between phylogenies and contact networks changes depending on epidemic dynamics and the fraction of infected hosts sampled. By considering pathogen phylogenies in a probabilistic framework, these coalescent models can also be used to estimate the statistical properties of contact networks directly from phylogenies using likelihood-based inference. We use this framework to explore how much information phylogenies retain about the underlying structure of contact networks and to infer the structure of a sexual contact network underlying a large HIV-1 sub-epidemic in Switzerland.

## Introduction

From the viewpoint of an infectious pathogen, host populations are highly structured by the physical contacts necessary for disease transmission to occur. For pathogens whose transmission does not require intimate or sustained physical contact, random mixing models assuming contacts form instantaneously between individuals may offer a reasonable approximation to the true dynamics of person-to-person contact. But for sexually-transmitted infections (STIs) and many other pathogens, the contacts required for transmission are generally more limited in number, less transient in nature, and form non-randomly based on individual behavior—resulting in host populations that are highly structured locally at the level of individuals [1–4]. For such pathogens, it is more reasonable to view communities as networks of individuals connected by edges that represent the physical contacts through which transmission can occur. Through the study of theoretical network models, epidemiologists now understand that contact network structure has a profound influence on epidemic dynamics and whether or not control strategies will be effective [5–9]. Yet studying the structure of contact networks empirically through methods such as contact tracing is difficult and costly, meaning we still know relatively little about real-world contact networks [10, 11].

New hope for the empirical study of contact networks has emerged in recent years from the widespread availability of pathogen molecular sequence data. In molecular epidemiology, sequence data is already commonly used to link individuals into probable transmission pairs or clusters based on the phylogenetic distances between their pathogens. While such approaches do not directly reveal the structure of contact networks, they can reveal paths in the contact network through which the pathogen spread and provide a useful heuristic for assessing how well connected networks are within and between different subpopulations or risk-groups [12–14]. Other methods in molecular epidemiology attempt to reconstruct the full details of the underlying transmission tree, the directed graph showing exactly who infected whom in an outbreak [15–20]. Essentially though, all current methods for inferring linkage and transmission trees take a bottom-up approach—they attempt to reconstruct routes of transmission by linking sampled individuals based on their phylogenetic distance. While this can be a powerful approach for studying densely sampled outbreaks where most infected individuals are sampled, bottom-up approaches may provide misleading results when applied to sparsely sampled epidemics. In this case, two infected individuals may have pathogens that are most closely related to one another phylogenetically but an unknown number of intervening infections might separate them in the true transmission tree. Thus the phylogenetic proximity of individuals may only weakly correlate with their proximity in the transmission tree, making it very difficult to reconstruct the detailed transmission history of who infected whom.

While it may not be possible to reconstruct the detailed structure of transmission networks from sparsely sampled data, it may still be possible to infer large-scale properties of contact networks. By simulating the phylogenetic history of pathogens spreading through networks, recent studies have shown that network properties can exert a strong influence on the structure of phylogenetic trees [21–23]. For example, increasing levels of contact heterogeneity—variation in the number of contacts individuals form—can result in increasingly asymmetric or imbalanced trees and shift the distribution of coalescent (i.e. branching) events earlier towards the beginning of an epidemic [21, 22]. However, statistical measures of tree topology like imbalance may only weakly correlate with network statistics like contact heterogeneity, and may be highly dependent on how samples are collected [23]. Moreover, in addition to network structure, population dynamics also strongly shape phylogenies and therefore potentially confound inferences of network structure drawn from phylogenies [23]. For example, clustering of samples together in phylogenetic trees has previously been assumed to indicate clustering of individuals in the underlying contact network, but phylogenetic clustering can arise naturally in epidemics even when no measurable degree of clustering exists in a population [24]. Taken together then, previous work suggests that contact network structure can shape pathogen phylogenies, but we do not yet know how to properly extract this information from trees.

In this paper, we present a new theoretical framework for relating pathogen phylogenies to contact networks using phylodynamic modeling. Our approach is quite different from bottom-up approaches in that it does not attempt to reconstruct the details of person-to-person transmission. Rather, we start with a random graph model [25] that captures the important statistical properties of real-world networks. We then use pairwise epidemic models [26–28] to capture the population dynamics of an epidemic on a network with the statistical properties specified by the random graph model. In addition to tracking the infectious status of individuals, these pairwise models track the status of pairs of individuals and thereby correlations in the infectious status of neighboring individuals, such as the depletion of susceptible hosts around infected individuals. Analogously, by shifting our focus from the level of individuals to the level of pairs, we derive a relatively simple coalescent model that captures a pathogen’s phylogenetic history as a backwards-time dynamical process on a network. The pairwise coalescent model naturally takes into account incomplete sampling and how network structure and epidemic dynamics interact to shape pathogen phylogenies. By considering phylogenies in a probabilistic framework, the pairwise coalescent model also allows us to compute the likelihood of a given phylogeny evolving on a network with defined statistical properties, and therefore to estimate the structure of networks from phylogenies using likelihood-based inference.

How local contact network structure shapes pathogen phylogenies has received some attention in recent years [21–23], but has not been comprehensively studied. After deriving the pairwise coalescent model, we therefore begin by using simulations to explore how network properties such as overall connectivity, clustering, contact heterogeneity and assortativity shape phylogenies. Using these simulations, we demonstrate that the pairwise coalescent model captures how these network properties shape phylogenies in terms of coalescent times, how lineages move through a network, and overall tree topology. We then go on to show that the model can be used to accurately estimate network properties from phylogenies, although how precisely depends strongly on sampling effort. Finally, we have implemented the model in BEAST 2 [29] as a package called PairTree, which we use to estimate the structure of a contact network underlying a large HIV sub-epidemic in Switzerland.

## Models and Methods

Our phylodynamic modeling framework is composed of three interacting components. The first two components, random graph and pairwise epidemic models, are well described in the literature and we only briefly review the necessary concepts and notation here. Instead, we focus on the third and novel component of our framework, the pairwise coalescent model, which we derive from the pairwise epidemic model.

### Random graph models

In network epidemiology, random graph models are often used to model the large-scale statistical properties of networks while treating the fine-scale details of who is connected to whom as random. Random graph models can therefore be thought of as a probability distribution on graphs constrained to take on certain statistical properties. Here, we use the configuration model [30] and extensions thereof to model network structure and generate random graphs parameterized to vary in overall connectivity, contact heterogeneity, clustering and assortative mixing.

#### A. Connectivity

Connectivity quantifies how well-connected individual nodes are in a network in terms of their degree, or number of contacts. In the simplest case of the configuration model, all *N* nodes are assigned a fixed degree 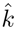 and then randomly connected to other nodes through edges, resulting in a homogenous or k-regular random graph. The parameter 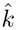 therefore quantifies the overall connectivity of the network.

#### B. Clustering

Clustering is defined as the probability that two nodes connected to a common neighbor are also connected to one another, and therefore quantifies how locally interconnected networks are [26]. Clustering can be quantified in terms of a clustering coefficient *ϕ*:

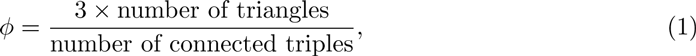

where a triangle refers to a closed loop of three connected nodes and a triple to three linearly connected nodes [25].

To introduce clustering into random networks, we use the triangular configuration model [31, 32]. Under this model, rather than defining the degree distribution *d_k_*, we define a joint degree distribution *d_st_* on the probability that a node is connected to *s* neighbors not forming triangles and 2*t* other neighbors through triangles, and thus has total degree *k = s* + 2*t.* As shown by [32], given *d_st_* and the overall degree distribution *d_k_,* the expected clustering coefficient is

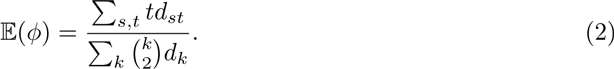

#### C. Contact heterogeneity

Contact heterogeneity refers to variation in the number of contacts individuals form in a network and can be quantified by the variance 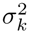 in the degree distribution *d_k_*. Networks with any arbitrary degree distribution can be generated under the standard configuration model by assigning each node *n* = 1, 2, …, *N* a degree according to a random degree sequence *k*_1_, *k*_2_, …, *k_N_* drawn from *d_k_*. Each individual node *n* is then randomly joined to *k_n_* other nodes to form the edges of the network.

#### D. Assortative mixing

Assortative mixing is the tendency for individuals to form connections with individuals similar to themselves, leading to correlations between the properties of adjacent nodes in a network [25]. Here, we consider assortative mixing by node degree. To introduce correlations in the degree of connected nodes, we specify the edge degree distribution *e_kl_,* which gives the probability of a randomly chosen edge connecting a degree *k* to a degree *l* node. The strength of assortative mixing can be quantified in terms of the assortativity coefficient *r* given *e_kl_* and *d_k_*:

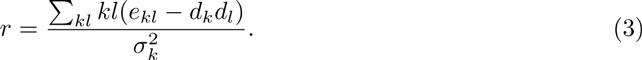

Thus, *r* can also be interpreted as the Pearson correlation coefficient in the degree of nodes connected by edges in the network.

To get a one-parameter random graph model that allows for the strength of assortative mixing to vary based on *r*, we follow [33] and constrain each entry in *e_kl_* to follow the form

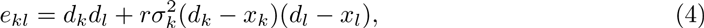

where *x_k_* is a normalized distribution chosen such that *e_kl_* is never negative. To our knowledge, there is no algorithm that allows for direct simulation of networks from the distribution over graphs defined by *e_kl_*. We therefore sample networks using the Metropolis-Hastings sampler also proposed by [33] that iteratively rewires networks until convergence on a target distribution defined by *e_kl_* is reached.

### Pairwise epidemic model

The second component of our modeling framework consists of epidemiological models that describe the dynamics of a pathogen spreading through a network with statistical properties specified by a random graph model. As in standard SIR-type epidemiological models, we track the infection status of each node or host as susceptible or infected, along with an optional recovered class. We use the notation [*S_k_*] and [*I_k_*] to denote the number of degree *k* susceptible and infected individuals; [*S_k_I_l_*] denotes the the number of pairs or edges in the network connecting *S_k_* and *I_l_* individuals. At the level of individuals, the epidemic dynamics are described by the following differential equations:

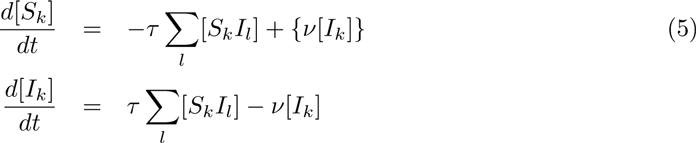

Here, *τ* is that rate at which infected individuals transmit to their neighbors and *ν* is the recovery rate. Terms in braces are either present if there is no immunity (the SIS model) or absent if infection is completly immunizing (the SIR model).

As seen from Eq (5), the transmission dynamics depend on the [*S_k_I_l_*] terms and thus how individuals are connected into pairs or partnerships. We therefore need to track the dynamics at the level of pairs:

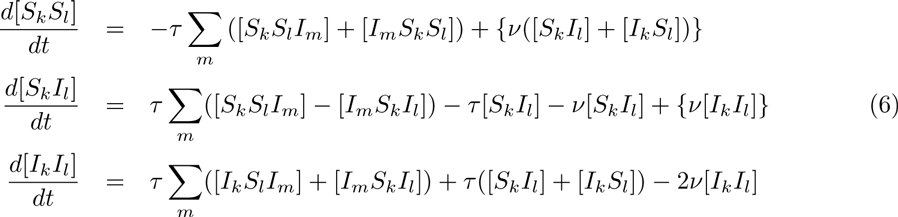

These are the pairwise network equations introduced by [26] and extended to heterogenous contact networks by [28]. By tracking the status of pairs rather than just individuals, the pairwise equations take into account local correlations that build up over time between the infection status of neighboring nodes; hence their other common name, correlation equations [27]. These local correlations arise because a node’s infection status depends strongly on the status of its neighbors. For example, early on in an epidemic positive correlations develop between infected individuals, reflecting the fact that infected individuals are likely to be surrounded by other infected individuals who either infected them or became infected by them. Because these correlations can have a strong impact on epidemic dynamics, such as through the local depletion of susceptible nodes surrounding infected nodes, tracking these correlations allows pairwise models to more accurately describe epidemic dynamics on networks.

While the dynamics at the level of pairs depends on the number of triples such as [*S_k_S_l_I_m_*], which in turn depends on even higher-order configurations, previous work has shown that moment closure methods can be used to approximate the number of triples based on the number of pairs without much loss of accuracy [28, 34]. We thus “close” the system at the level of pairs by approximating each triple of arbitrary type [*ABC*] as:

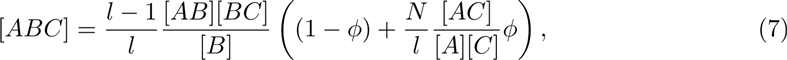

where *l* is the degree of the central node in state *B.* By taking into account the clustering coefficient *ϕ*, this moment closure takes into account additional state correlations that can arise between three nodes when there is appreciable clustering in the network [26, 27]

### Pairwise coalescent model

The third and novel component of our modeling framework are coalescent models that allow us to probabilistically relate the phylogenetic history of a pathogen back to the dynamics of an epidemic on a network. In essence, these coalescent models provide a probability distribution over trees on random networks, and therefore allow us to compute the likelihood of a given phylogeny having evolved on a network. While coalescent theory has previously been extended to accommodate the nonlinear transmission dynamics of infectious pathogens [35–38], these coalescent models assume random mixing, at least within discrete subpopulations, and therefore neglect local contact network structure. Below, we extend the structured coalescent models of [38] to include local contact network structure by shifting our focus from the level of individuals to pairs of hosts in the network.

The likelihood of a phylogeny 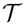 under a structured coalescent model with parameters *θ* has the general form:

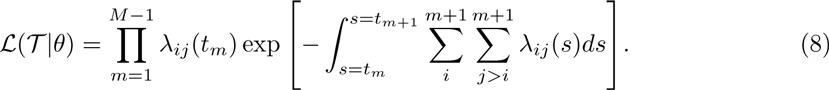

For a tree containing *M* samples, the total likelihood is the product of the likelihood of each of the *M* – 1 coalescent events and the waiting times between events. The likelihood of each coalescent event is given by the rate *λ_ij_*(*t_m_*) at which lineages *i* and *j* coalesce at the time of the event *t_m_.*

The pairwise coalescent rates *λ_i__j_* are centrally important to our model as they are required to compute the likelihood in Eq (8) and provide the main link between the epidemic dynamics and the coalescent process. To derive these rates, we begin by making the simplifying assumption common in phylodynamics that only a single pathogen lineage resides in each infected host. While this assumption ignores within-host pathogen diversity, it dramatically simplifies the relationship between transmission events and coalescent events in the pathogen phylogeny: each coalescent event in the phylogeny will represent a transmission event on the network. Below, we use this relationship to derive the pairwise coalescent rates *λ_ij_* for pairs of lineages.

#### Pairwise coalescent rates

To derive the pairwise coalescent rate *λ_ij_*, we first need to consider the probability that lineages *i* and *j* coalesce conditional on a transmission event occurring somewhere in the network. In order for two lineages to coalesce at a transmission event from an individual with *l* contacts to an individual with *k* contacts, we can reason that at the time of the event three conditions must hold:

1. The two lineages must be in two infected individuals, one with *k* and the other with *l* contacts.
2. The two lineages must reside in two nodes connected in a *I_k_I_l_* pair.
3. The two lineages must be in the specific *I_k_I_l_* pair involved in the transmission event.

We note that each of these conditions must be met in turn for the remaining conditions to be met. We will therefore consider the probability that each of these conditions is true in turn conditional on the preceding conditions having been met.

First, consider the probability that lineages *i* and *j* are in two infected individuals; one in a *I_k_* node and the other in a *I_l_* node. In general, we will not know the degree of the infected node in which the lineage resides (for shorthand, we will refer to this as the lineage’s state). We must therefore treat the state of lineages probabilistically and will use the notation *p*_ik_ to represent the probability that lineage *i* resides in a degree *k* infected node. The probability 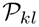 that lineages *i* and *j* reside in nodes with degrees *k* and *l* is then equal to the probability that lineage *i* is in state *k* and lineage *j* is in state *l* or vice versa, such that

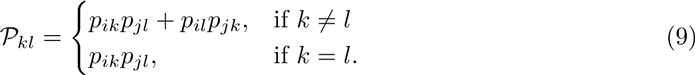

Second, consider the probability that lineages *i* and *j* reside in two nodes connected in a *I_k_I_l_* pair. The total number of possible pairs between *I_k_* and *I_l_* nodes is [*I_k_*] [*I_l_*] if *k* ≠ *l* or 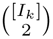 if *k* = *l.* Because pairs are assumed to form randomly under our random graph models, the probability *χ_kl_* that a randomly chosen *I_k_* node is connected to a random *I_l_* node in a *I_k_I_l_* pair is:

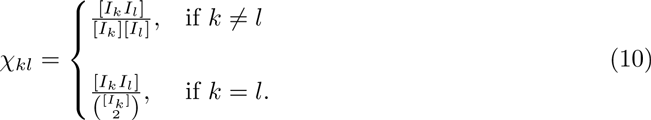

Given that our two lineages reside in *I_k_* and *I_l_* nodes, then the probability that our lineages reside in a *I_k_I_l_* pair is *χ_kl_.*

Third, given that lineages *i* and *j* are in two nodes that form a *I_k_I_l_* pair, the probability that it is this pair out of all *I_k_I_l_* pairs in the network that was involved in a given transmission event is simply 1/[*I_k_I_l_*].

These three probabilities collectively give the probability that lineages *i* and *j* coalesce at a particular transmission event. Multiplying these probabilities by the total rate at which degree *l* nodes transmit to degree *k* nodes, the rate at which lineages *i* and *j* coalesce through *l* → *k* transmission events is:

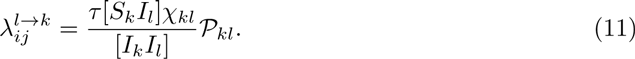

Summing over all possible transmission events with respect to the degree of the nodes involved, we arrive at the total pairwise coalescent rate:

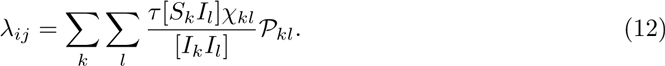

The likelihood given in Eq (8) can then be computed by plugging in *λ_ij_*(*t*) for each lineage pair after numerically integrating the ODEs for [*S_k_I_l_*] and [*I_k_I_l_*] given in Eq (6) up to time *t.*

For a homogenous network where all nodes have the same degree 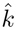 Eq (12) simplifies to

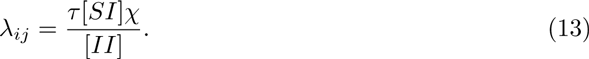

In a fully connected network where each node has degree 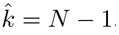, every node is connected to every other node. In this limiting case, we expect the dynamics of an epidemic on a network to be the same as under a random mixing model with a transmission rate *β* scaled so that infectious contacts occur at the same rate under both models. In this case, the probability that two random infected nodes are connected in a pair 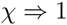, 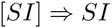, and 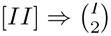. Making these substitutions in Eq (13), we see that

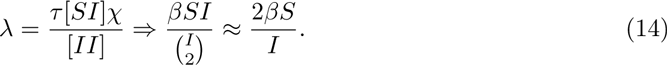

This is the same pairwise coalescent rate derived by [35] for a random mixing SI(R) model. We therefore see that the pairwise coalescent model and earlier coalescent models assuming random mixing converge in the limit of a fully connected network.

#### Tracking lineage movement

We now consider how individual pathogen lineages move through a network. Because we need to know the probabilities *p_ik_* of a lineage residing in a degree *k* host in order to compute the pairwise coalescent rates given in Eq (12), we probabilistically track the movement of lineages by tracking how *p_ik_* changes backwards through time along a lineage using a framework based on master equations previously developed by [38]. These master equations have the general form

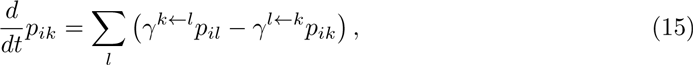

where *γ^k←l^* is the rate at which lineages transition from degree *l* to degree *k* hosts backwards in time. How these transition rates are computed and further details about how the ancestral degree distribution *p_k_* is computed for each lineage in a phylogeny are described in the SI Text, where we also show that these master equations accurately describe how lineages move through networks.

## Results

### The coalescent process on random networks

Most of the local structure present within real-world contact networks can be captured by a few key statistical properties: its overall connectivity and the degree of clustering, contact heterogeneity and assortativity [28]. To see how these network properties shape pathogen phylogenies and how well the pairwise coalescent model captures their effects, we generated networks under random graph models parameterized to obtain networks with known statistical properties. On top of these networks, we simulated the spread of an epidemic using individual-based stochastic (IBS) simulations that tracked the ancestry of each pathogen lineage forward in time so that a true phylogeny was obtained from each simulation (see SI Text). We then compared the epidemic dynamics and phylogenies simulated under the the IBS model to those expected under the pairwise epidemic and coalescent models.

As expected from earlier work [9, 34, 39], the pairwise epidemic model provides an excellent deterministic approximation to the mean dynamics observed in IBS simulations across a wide range of random networks, whereas random mixing models generally do not (Fig 1). Likewise, the pairwise coalescent model does an excellent job of capturing the coalescent process on these networks in terms of the temporal distribution of coalescent events over the epidemic (Fig 2). In contrast, the coalescent distributions expected under a random mixing coalescent model provide a reasonable approximation on some networks but not others (Fig 2). For example, on poorly connected and highly clustered networks, the expected distribution of coalescent times under random mixing deviates widely from the IBS simulations. This is the case even if we condition the random mixing model on the more accurate population trajectories predicted by the pairwise epidemic model. On better connected networks and on networks with more contact heterogeneity, the random mixing model does almost as well as the pairwise coalescent model. In the SI Text, we additionally explore when the pairwise approximation fails due to the presence of higher-order network structure.

**Fig 1.**
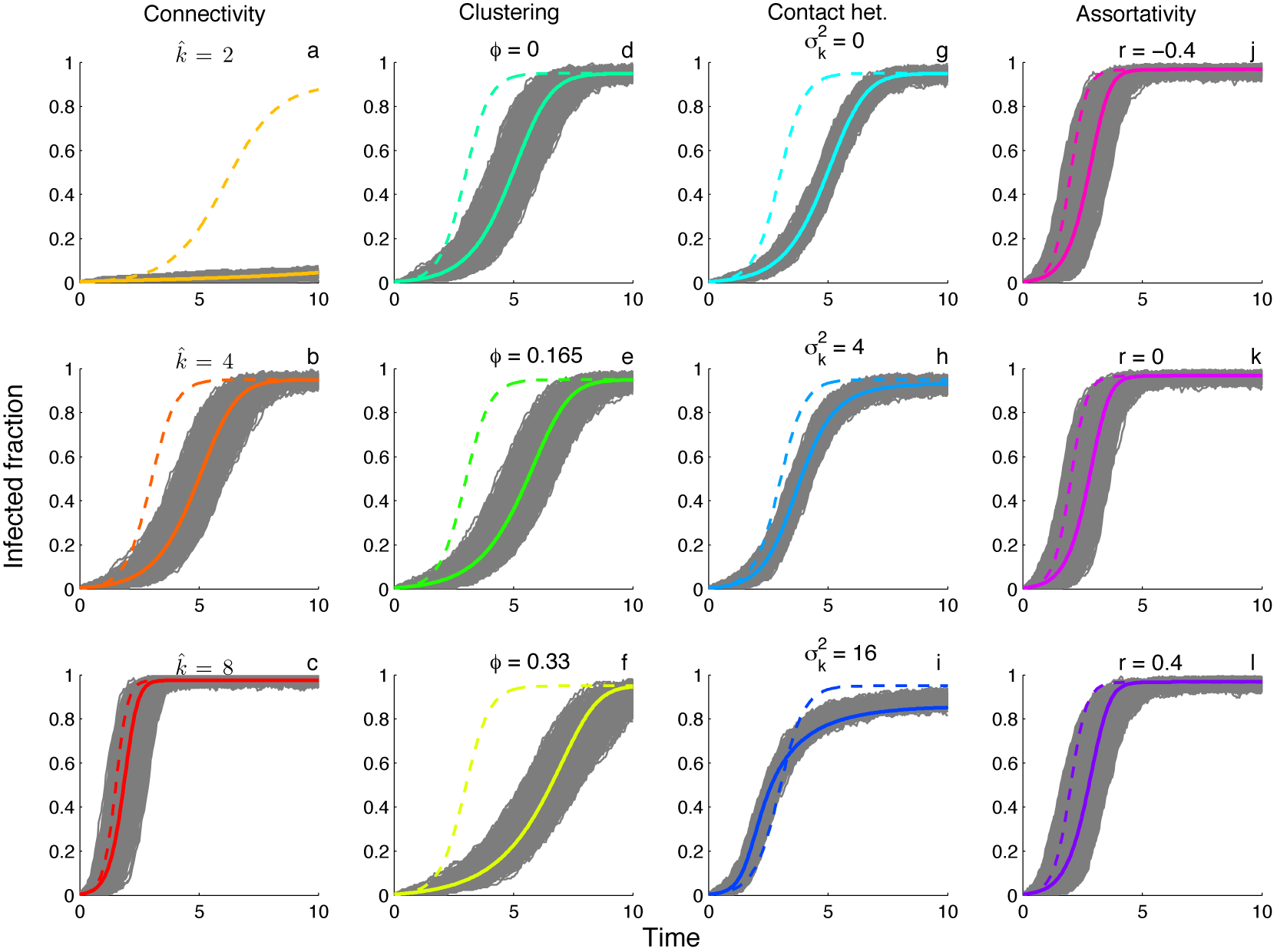
Comparison of SIS epidemic dynamics on networks with different statistical properties. Grey lines show 500 stochastic realizations of the individual-based model run on different random networks. Colored lines show the mean dynamics expected under the pairwise epidemic model (solid) versus the random mixing model (dashed). **(a-c)** Homogenous networks with varying overall connectivity 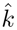. **(d-f)** Homogenous networks with different clustering coefficients *ϕ* but fixed degree 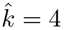. **(g-i)** Heterogenous networks with constant mean degree *μ_k_* = 4 but with different variances 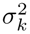 in the degree distribution. **(j-l)** Heterogenous networks with different assortativity coefficients *r*. To allow for greater assortativity, the mean and variance of the degree distribution was raised to six. For all simulations the network size *N* = 250, the transmission rate *τ* = 0.5 and the recovery rate *v* = 0.1. The transmission rate *β* under random mixing was scaled so that the rate of infectious contacts was the same as under the pairwise epidemic model at *t* = 0, giving the two models the same intrinsic growth rate.

**Fig 2.**
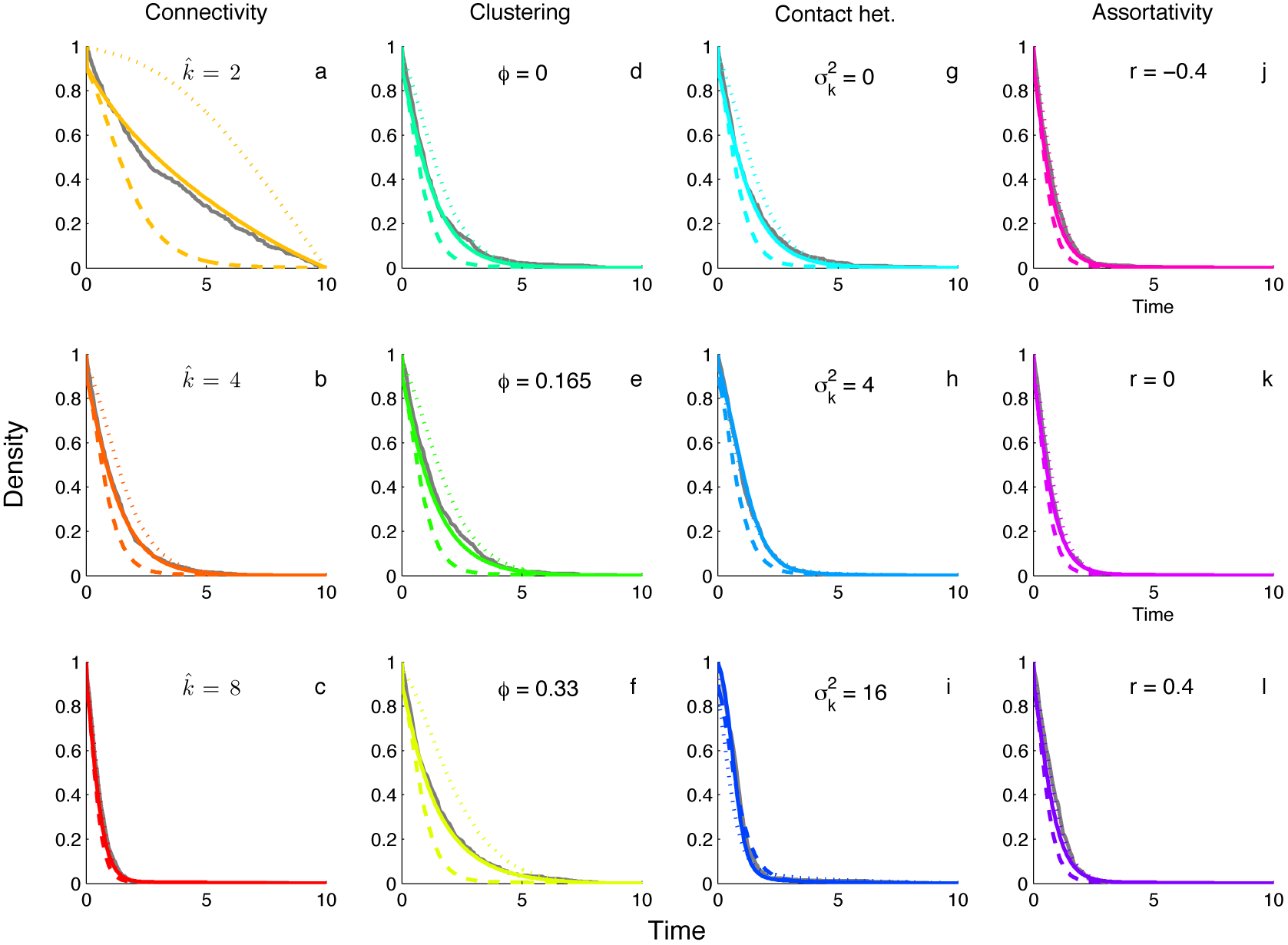
The cumulative distribution of coalescent event times for the networks and epidemic dynamics shown in Figure 1. Grey lines show the observed distribution of coalescent events obtained by tracking two randomly sampled lineages back in time until they coalesce in 500 stochastic individual-based simulations. Colored lines show the theoretically predicted coalescent densities under the pairwise coalescent (solid), the random mixing coalescent (dashed) and the random mixing coalescent conditioned on the mean dynamics provided by the pairwise epidemic model (dotted). **(a-c)** Homogenous networks with varying overall connectivity 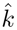. **(d-f)** Homogenous networks with different clustering coefficients *ϕ*. **(g-i)** Heterogenous networks with different variances 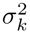 in the degree distribution. **(j-l)** Heterogenous networks with different assortativity coefficients *r*. All network and epidemiological parameters are the same as in Figure 1.

### Network effects on tree topology

In addition to coalescent time distributions, contact network structure can shape the topology of phylogenies. In particular, trees tend to become increasingly asymmetric or imbalanced as the amount of contact heterogeneity increases in a population [21–23, 40]. We therefore simulated trees on random networks with different levels of contact heterogeneity using IBS simulations and under the pairwise coalescent model using backward-time simulations in order to see if the coalescent model can capture the effects of contact heterogeneity on tree imbalance.

Overall, trees simulated under the pairwise coalescent model are very similar in shape to trees simulated on random networks using IBS simulations (Fig 3). For two different measures of imbalance, Colless’ and Sackin’s index, tree imbalance grows only weakly with increasing contact heterogeneity for both coalescent trees and IBS trees. While imbalance in our coalescent trees grows proportionally to how imbalance grows in IBS trees with increasing contact heterogeneity, coalescent trees do tend to be slightly more imbalanced, especially for Sackin’s index. The number of cherries, or pairs of tips sharing a direct ancestor, can also be used as a measure of imbalance as more imbalanced trees will have fewer cherries [40]. We found that the number of cherries decreases proportionally for both coalescent and IBS trees as contact heterogeneity increases, although again coalescent trees seem to be slightly more imbalanced with fewer overall cherries (Fig 3). Thus, it appears that the pairwise coalescent model can capture the effects of local contact structure on tree topology, even if these effects are rather weak overall.

**Fig 3.**
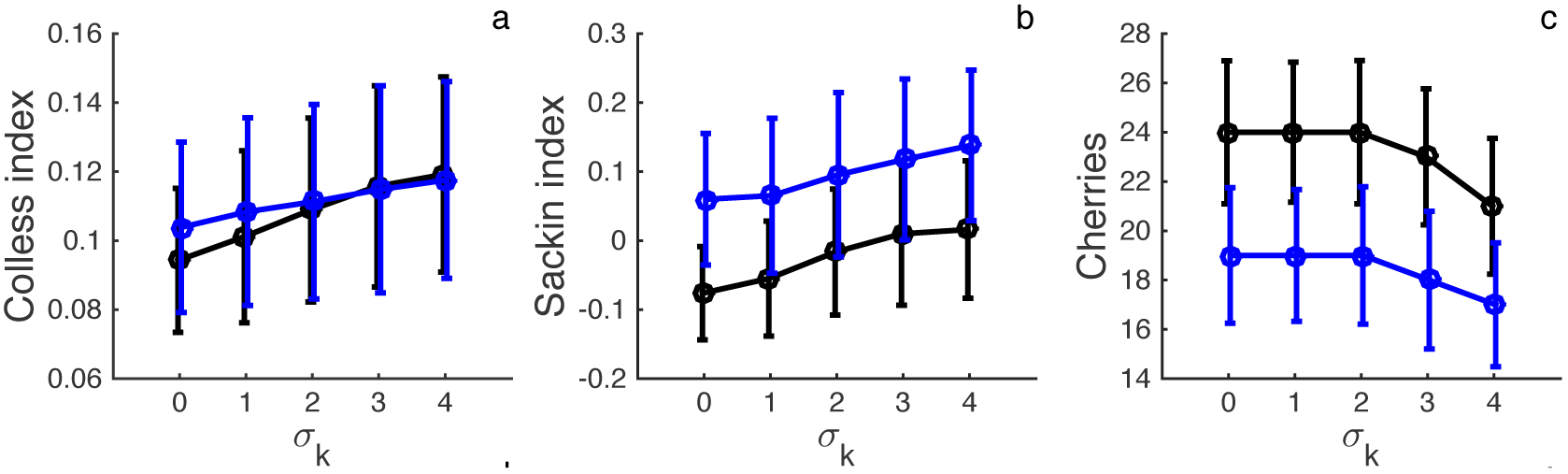
Phylogenetic tree imbalance on networks with increasing levels of contact heterogeneity. Trees were simulated using stochastic, individual-based simulations on random networks (black lines) or using backward-time simulations of the pairwise coalescent model (blue line). **(a-c)** Imbalance measured in terms of Colless’ index, Sackin’s index and the total number of cherries for trees simulated with increasing levels of contact heterogeneity. Colless’ and Sackin’s imbalance measures are normalized according to the expected imbalance of a tree with the same number of samples. Circles and vertical lines mark the mean and standard deviation of the imbalance measures for 500 simulated trees. All simulations were performed with *N* = 250 and a sampling fraction of *ρ* = 0.5.

### Inference

The pairwise coalescent model allows us to compute the likelihood of a given phylogeny being generated by an epidemic on a random network with defined statistical properties. It is therefore possible to directly estimate the statistical properties of a network directly from a phylogeny. However, phylogenies may retain little information about contact network structure, especially if the epidemic is sparsely sampled. To explore the information content of phylogenies regarding network structure, we simulated epidemics on random networks with known statistical properties. A variable fraction of infected nodes was then sampled upon removal to obtain phylogenies with sampling fractions *ρ* of 10, 25, 50 and 100%. The pairwise coalescent model was then used to construct likelihood profiles for different parameters controlling local network structure. Here, we focus only on the information content of the phylogenies but we further investigate the statistical performance of the pairwise coalescent model as an estimator of these parameters in the SI Text.

At sampling fractions at or below 10%, except for overall connectivity the simulated phylogenies contain little or no information about local network structure, as seen from the essentially flat likelihood profiles (Fig 4). At sampling fractions ≥ 25%, the likelihood profiles begin to show significant curvature for clustering and contact heterogeneity, and with sampling fractions ≥ 50% the likelihood profiles are sharply curved enough that these parameters can be estimated rather precisely with narrow 95% confidence intervals. Assortativity appears more difficult to infer from phylogenies, even if the true degree of sampled nodes is provided (Fig 4). Although the likelihood profiles for *r* do show some curvature at sampling fractions ≥ 50%, the credible intervals remain relatively wide even with complete sampling.

**Fig 4.**
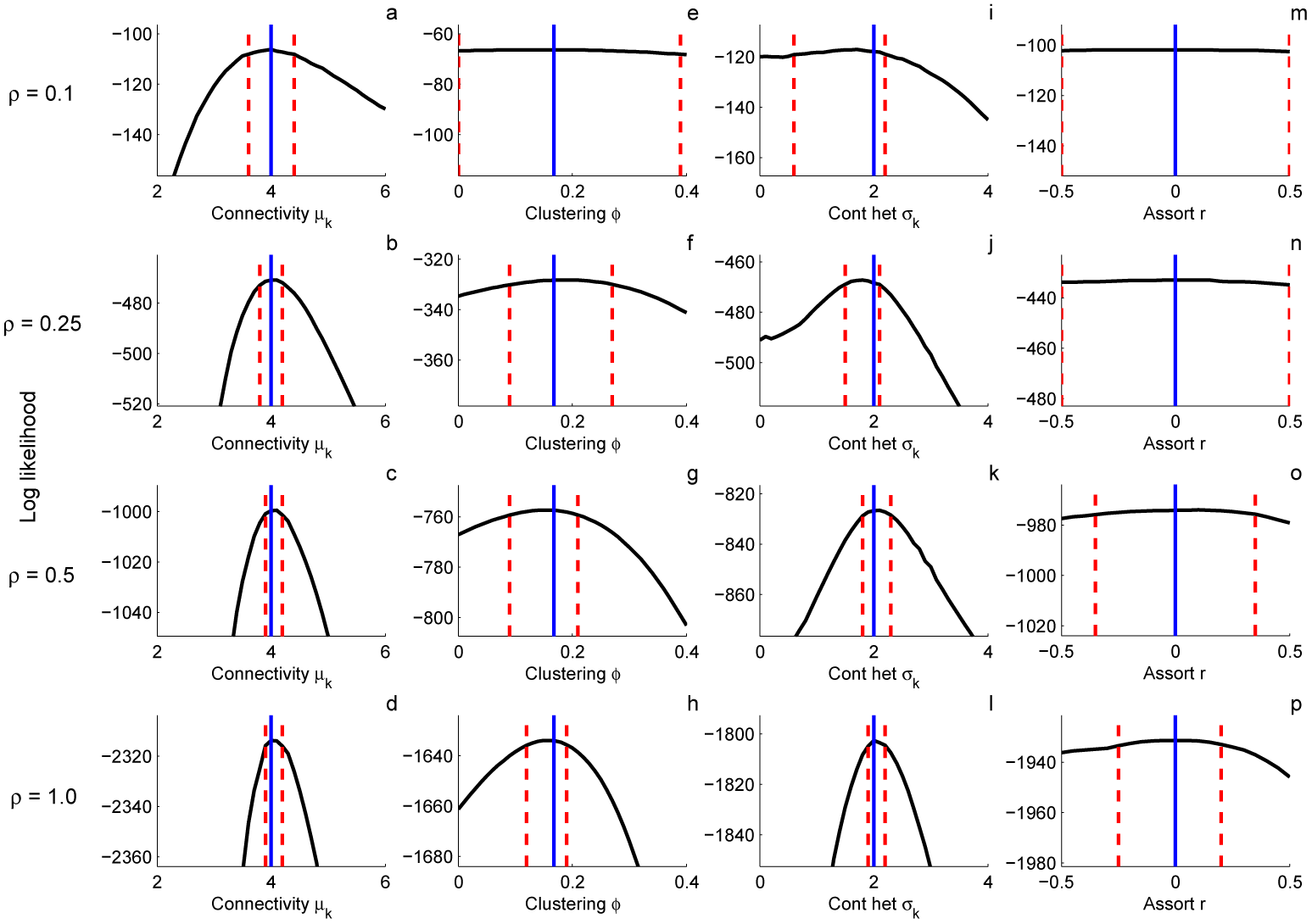
Likelihood profiles for network parameters inferred from phylogenies containing a variable fraction of all infected nodes *ρ*. Solid blue lines indicate true parameter values and dashed red lines are the approximate 95% confidence intervals derived from the quantiles of the chi-squared distribution with one degree of freedom. Parameters were estimated one at a time, with all other parameters fixed at their true values. **(a-d)** Overall connectivity as parameterized by the mean degree *μ_k_*. Here we assume the degree distribution follows a discretized gamma distribution with a fixed standard deviation *σ_k_* = 2 **(e-h)** Clustering as parameterized by the clustering coefficient ϕ. **(i-l)** Contact heterogeneity as parameterized by the standard deviation of the degree distribution *σ_k_*. The mean of the degree distribution was fixed at *μ_k_* = 4. **(m-p)** Assortativity as parameterized by the assortativity coefficient r of the edge degree distribution. For assortativity, the true degree of all sampled nodes was provided when computing the likelihood profiles. Trees were simulated on networks with *N* = 250 and the epidemiological parameters were fixed at the values used in Figures 1 and 2.

### HIV-1 in Switzerland

We performed a phylodynamic analysis of a HIV-1 subtype B epidemic among men-who-have-sex-with-men (MSM) in Switzerland using the pairwise coalescent model to see if we could estimate the statistical properties of a real-world sexual contact network. HIV *pol* sequences from infected individuals were obtained from patients enrolled in the Swiss HIV Cohort Study [41–43]. To minimize the effects of spatial structure, we focus on a single large sub-epidemic identified as primarily occurring in the Zürich region in a preliminary phylogenetic analysis (see SI Text). A time-calibrated phylogeny containing 200 sequences revealed this sub-epidemic to be quite genetically diverse with many older lineages originating in the early 1980’s (Fig 5).

**Fig 5.**
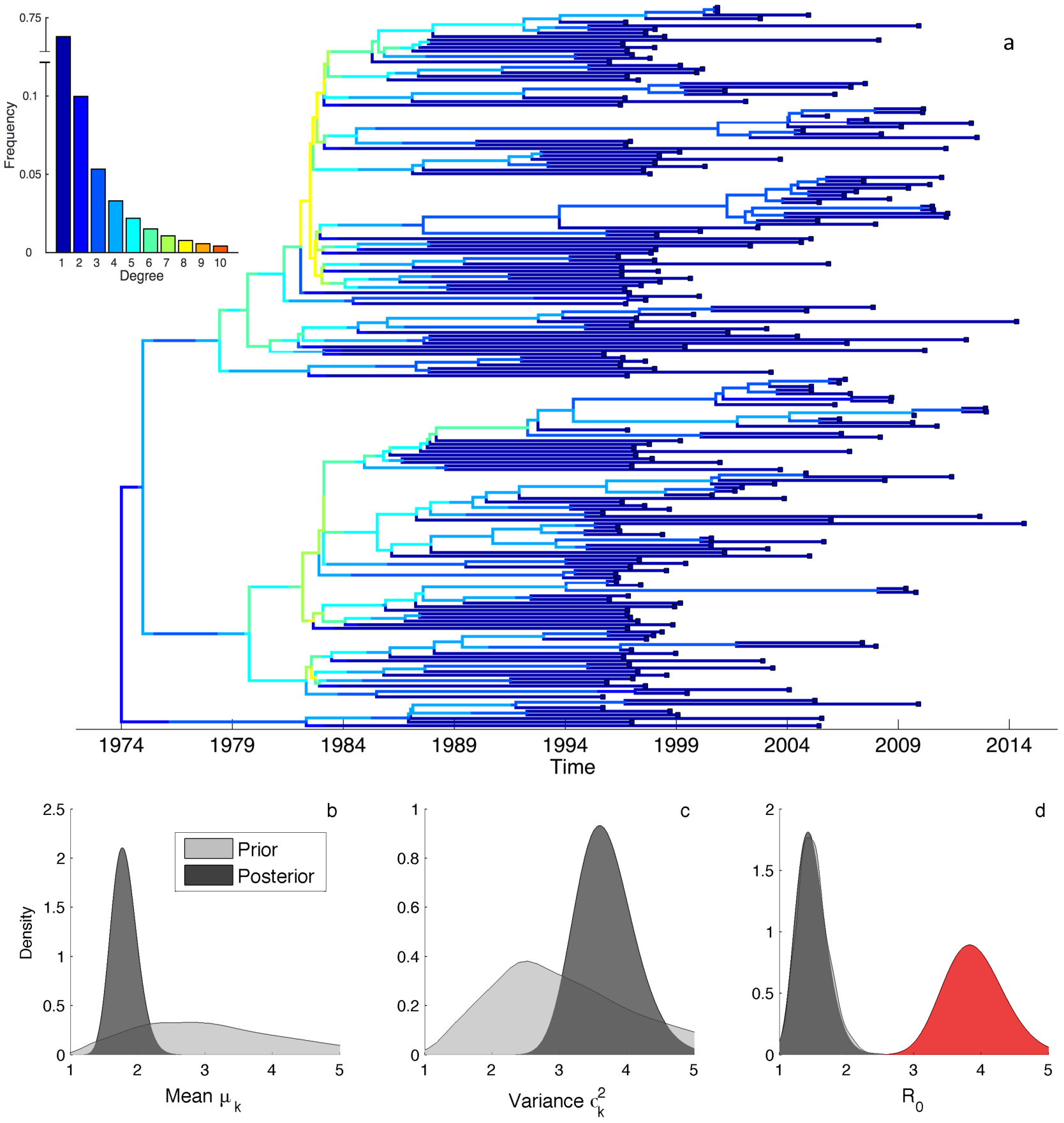
Phylodynamic analysis of a HIV-1 sub-epidemic among MSM in Switzerland. **(a)** Maximum clade credible phylogeny reconstructed from 200 HIV *pol* sequences. Lineages are colored by their expected degree in the network based on their ancestral degree distribution inferred under the pairwise coalescent model. The ancestral degree distribution was computed using a forward-backwards type algorithm while conditioning on the median posterior estimate of all parameters. Inset shows the inferred degree distribution for the entire network (capped at *k* = 10 for visual simplicity), (**b-c**) Posterior (dark grey) and prior (light grey) distributions of the mean degree *μ_k_* and variance 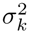 of the degree distribution, **(d)** Estimates of the basic reproductive number *R*_0_ inferred under the pairwise coalescent (grey) and under the random mixing coalescent model (red).

We fit a SIR-type pairwise epidemic model to the dated phylogeny assuming a discretized gamma distribution for the degree distribution *d_k_*. This allowed us to independently estimate the mean *μ_k_* and variance 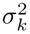 of the network’s underlying degree distribution. The posterior estimates of *μ_k_* and 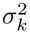 indicate that the network was not especially well-connected (median *μ_k_* = 1.78) but heterogenous in degree (median 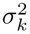 = 3.65) (Fig 5). The basic reproductive number was estimated to be between 1.0 and 2.0, although unlike for the network parameters the posterior density of *R*_0_ did not diverge appreciably from the prior (Fig 5). *R*_0_ values estimated under the pairwise coalescent model were however significantly lower than the values estimated under the random mixing model. Because the random mixing model does not account for contact heterogeneity, it can only capture the rapid early growth of the epidemic by overestimating *R*_0_.

Overall our phylodynamic analysis suggests that this particular sub-epidemic spread rapidly by way of a few highly connected individuals. This is supported by the inferred degree distribution of the network and can be seen from the expected degree of lineages computed from the inferred ancestral degree distribution of each lineage over time (Fig 5). Most coalescent (i.e. transmission) events early in the epidemic are attributable to lineages residing in high degree individuals. Later, towards the beginning of the 2000’s, a few clusters in the tree begin to grow again through new transmission events along lineages with higher than average degree, which corresponds in time to the resurgence of HIV among MSM in Switzerland [42, 44].

## Discussion

Recent work has suggested that the structure of local contact networks can shape pathogen phylogenies [21–23]. Yet it remains unclear how much information pathogen phylogenies retain about the networks through which they spread and how to best extract information about network structure from trees. As a step towards addressing these questions, we sought a simple theoretical framework to explore the relationship between contact networks, epidemic dynamics, and phylogenies. Starting with random graph and pairwise epidemic models, we derived a fairly simple coalescent model that includes local network structure by using a pair approximation technique. By treating the coalescent process as a backwards-time dynamical process on a network, our pairwise coalescent model allows us to capture the phylogenetic history of a pathogen in terms of how lineages move through a network and the rates at which they coalesce. As we have shown, our phylodynamic modeling framework provides a very good approximation to the coalescent process on random networks and can recapitulate the major features of pathogen phylogenies simulated on different types of random graphs.

Using the pairwise coalescent model and individual-based stochastic simulations as guides, we reexamined how contact network structure shapes pathogen phylogenies. Overall, we found that local contact network structure can have a strong impact on the the coalescent process in terms of the timing of coalescent events. Network properties like overall connectivity and contact heterogeneity that increase the epidemic growth rate concentrate coalescent events towards the beginning of an epidemic, while properties like clustering that slow epidemic growth broaden the distribution of coalescent events over the epidemic. On the other hand, properties like assortativity that have no strong effect on epidemic dynamics likewise have little influence on the timing of coalescent events. This suggests that local contact network structure primarily shapes the coalescent process indirectly through the network’s influence on epidemic dynamics, particularly the timing of transmission events. The timing of transmission events as regulated by local interactions on the network also appears to determine how well random mixing models can approximate the coalescent process on networks. On weakly connected or highly clustered networks where local interactions strongly limit transmission due to saturation effects, random mixing models overshoot the true transmission rate and therefore also the expected coalescent rate. In better connected networks, the effect of these local interactions on transmission is minimized by well-connected nodes and the random mixing models can perform quite well. Local contact network structure can therefore probably be safely ignored in highly connected networks, but may be important to consider in less well-connected networks.

Because the pairwise coalescent model can be used for likelihood-based inference, it offers a means of exploring how much information phylogenies contain about contact network structure. Using simulated phylogenies, we found that our ability to infer network properties was highly dependent on the fraction of sampled individuals. We could estimate network properties that strongly regulate epidemic dynamics, such as overall connectivity and the degree of contact heterogeneity, even at sampling fractions as low as 10-25%. Other properties that do not strongly regulate epidemic dynamics, such as assortativity, proved difficult to precisely estimate even with complete sampling. This observation suggests that for the parameters that can be estimated at low sampling fractions, we may largely be inferring the structure of networks not from any direct signal of network structure in the tree itself, but from the indirect effect of network structure on the epidemic dynamics reflected in the timing of the coalescent events in the tree.

While it therefore appears difficult to estimate some network properties from phylogenies, we were able to estimate the degree distribution of a sexual contact network underlying a large HIV sub-epidemic in Switzerland. While we were likely helped by the high fraction of HIV infected individuals sampled in Switzerland and the relatively informative priors we placed on the model’s epidemiological parameters, this demonstrates that it is at least technically possible to estimate the structure of real-world contact networks from phylogenies. Our analysis of the Swiss HIV data also indicated that accounting for network structure in phylodynamic models can be important for estimating key epidemiological parameters. Our estimated values of the net reproductive number *R*_0_ under a model assuming random mixing were more than twice as high as under a pairwise coalescent model that allowed for contact heterogeneity. Phylodynamic methods based on random mixing models may therefore be inappropriate when host populations are highly locally structured or when contact patterns vary considerably among individuals.

While we strove for simplicity, the true complexity of real-world contact networks does highlight some deficiencies in the pairwise models. First, while we only considered perfectly static random graph models, real-world networks temporally evolve as new contacts form and dissolve. Pairwise epidemic models that allow for dynamic partner exchange have been proposed [45, 46], and in theory could be merged with our pairwise coalescent model to explore contact durations that are intermediate between the infinitesimal nature assumed by random mixing models and the permanent nature assumed by static models. Finally, the random graph models we employed here only consider local structure at the level of pairs in the network. Higher-order structure that subdivides networks into different communities also likely plays a very strong role in shaping pathogen phylogenies. Developing methods that can quantify connectivity within and between communities while accounting for epidemic dynamics and incomplete sampling such as our approach does on local networks remains a challenging but highly important area of future research.

## Acknowledgments

We would like to thank Dr. Gabriel Leventhal for his insightful comments on this work. DAR is funded by the ETH Zürich Postdoctoral Fellowship Program and the Marie Curie Actions for People COFUND Program. RK is supported by Swiss National Science Foundation (SNF, grant BSSGI0_155851). TS is supported in part by the European Research Council under the Seventh Framework Programme of the European Commission (PhyPD: grant agreement number 335529). In addition, the Swiss HIV Cohort and the SHCS resistance database were supported by the SNF (grant 33CS30_148522, grant 320030 159868 to HFG, and grant PZ00P3-142411 to RK); the Swiss HIV Cohort Study (SHCS; projects 470, 528, 569, and 683); the SHCS Research Foundation; the Yvonne-Jacob Foundation; Gilead, Switzerland (one unrestricted grant to the SHCS Research Foundation and one unrestricted grant to HFG); and the University of Zürich’s Clinical Research Priority Program (Viral infectious diseases: Zürich Primary HIV Infection Study; to HFG). We thank the patients who participate in the SHCS; the physicians and study nurses, for excellent patient care; the resistance laboratories, for high-quality genotypic drug resistance testing; SmartGene (Zug, Switzerland), for technical support; Brigitte Remy, RN, Martin Rickenbach, MD, Franziska Schöni-Affolter, MD, and Yannick Vallet, MSc, from the SHCS Data Center (Lausanne, Switzerland), for data management; and Dani`ele Perraudin and Mirjam Minichiello, for administrative assistance. The members of the SHCS are Aubert V, Battegay M, Bernasconi E, Böni J, Braun DL, Bucher HC, Burton-Jeangros C, Calmy A, Cavassini M, Dollenmaier G, Egger M, Elzi L, Fehr J, Fellay J, Furrer H (Chairman of the Clinical and Laboratory Committee), Fux CA, Gorgievski M, Günthard H (President of the SHCS), Haerry D (deputy of “Positive Council”), Hasse B, Hirsch HH, Hoffmann M, Hösli I, Kahlert C, Kaiser L, Keiser O, Klimkait T, Kouyos R, Kovari H, Ledergerber B, Martinetti G, Martinez de Tejada B, Marzolini C, Metzner K, Müller N, Nadal D, Nicca D, Pantaleo G, Rauch A (Chairman of the Scientific Board), Regenass S, Rudin C (Chairman of the Mother and Child Substudy), Scherrer A, (Head of Data Centre), Schmid P, Speck R, Stöckle M, Tarr P, Trkola A, Vernazza P, Weber R, Yerly S.

## Supporting Information

### Tracking lineage movement on networks

Here we consider how lineages move through a network in terms of the ancestral degree distribution of a lineage. Going backwards in time, the degree of a lineage will transition from *k* to *l* whenever the lineage is transmitted from a degree *l* individual to a degree *k* individual in forward time. Transitions from *k* to *l* in backwards time are written as *l ← k* so that the direction of time is transparent. With incomplete sampling, a lineage may be transmitted between two nodes at a coalescent event that went unobserved in the tree because the parent lineage was not sampled. A lineage will therefore transition between states along branches in the tree each time an unobserved transmission event occurs between nodes of unequal degree. Thus, the rate at which *l ← k* transitions occur along a lineage currently in state *k* is equal to the rate at which the lineage coalesces with lineages in state *l* (through a *l → k* transmission event) that are not among the sampled lineages in the phylogeny. Assuming for the moment that there are no lineages in the phylogeny currently in state *l*, the rate at which *l ← k* transitions occur along a branch is

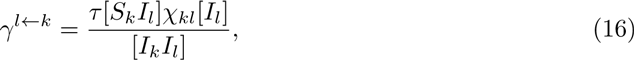

which is the rate of coalescence between a lineage in state *k* and all [*I_l_*] lineages in the population.

Notice that if *k ≠ l,* then *χ_kl_* = [*I_k_I_l_*]/[*I_k_*]][*I_l_*], Eq (16) simplifies to

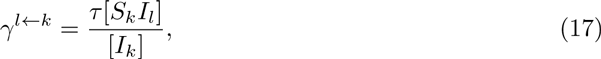

which has the more intuitive interpretation that a lineage transitions from state *k* to *l* at the same rate at which *l* → *k* transmission events occur in the population multiplied by the probability 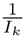 that it is this particular lineage in state *k* that is transmitted.

If the phylogeny contains lineages in state *l*, we need to consider that in order for the coalescent event to appear as *l* ← *k* transition along a branch, the parent lineage must not be among the sampled lineages in the phylogeny. As suggested by [38], the expected number of sampled lineages al in state *l* can be approximated from the lineage state probabilities as 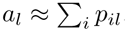. We can then substitute the [*I_l_*] term in Eq (16) with the probable number of lineages in state *l* but not in the phylogeny: [*I_l_*] — *a_l_.*

Given these transition rates, we can write down master equations for how *p_ik_* changes backwards in time:

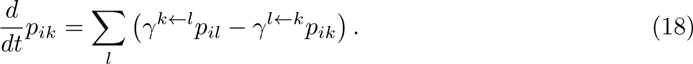

These master equations allow us to compute the probability of a lineage being in a given state at any time in the past, which we refer to as the ancestral degree distribution of a lineage. However, we generally do not know the degree of sampled individuals, we need to place a prior on *p_ik_* at the time of sampling. We use the degree distribution of the infected population at the time of sampling as a natural prior on the initial values of *p_ik_*:

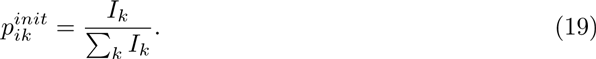

To obtain the degree distribution of the infected population, we can numerically solve the ODEs for *I_k_* under the pairwise epidemic model.

Finally, we need to consider how the lineage state probabilities get updated after an observed coalescent event in the tree. Specifically, we need to compute the state probabilities for the parent lineage *h* after its daughter lineages *i* and *j* coalesce. This is:

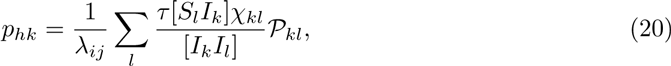

which is just the normalized probability of the parent being in state *k* conditional on 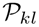 and the rate at which transmission events occur.

In order to see if the master equations given in Eq (18) provide an accurate representation of how lineages move through a network, we compare our theoretical expectations of *p_k_* with stochastic simulations where we recorded the state of a single sampled lineage backwards through time in each realization. In the population at large, well-connected nodes with high degree are overrepresented in the infected population early in an epidemic but the degree distribution of infected nodes rapidly converges to a stationary and approximately uniform distribution where all nodes have an equal probability of being infected regardless of degree (SI Fig 1a). Relative to the infected population at large, the ancestral degree distribution reconstructed from IBS simulations reveals that sampled lineages have an even higher probability of being in well-connected nodes during the early stages of an epidemic (SI Fig 1b). This results from lineages in higher degree nodes leaving more descendants and therefore having a higher probability of being ancestral to a sampled lineage. The master equations used by the pairwise coalescent model to track lineage movement reproduce this pattern almost exactly, although there is some disagreement during the earliest stages of the epidemic when *I_k_* << 1 for all *k* (SI Fig 1c).

**SI Figure 1.**
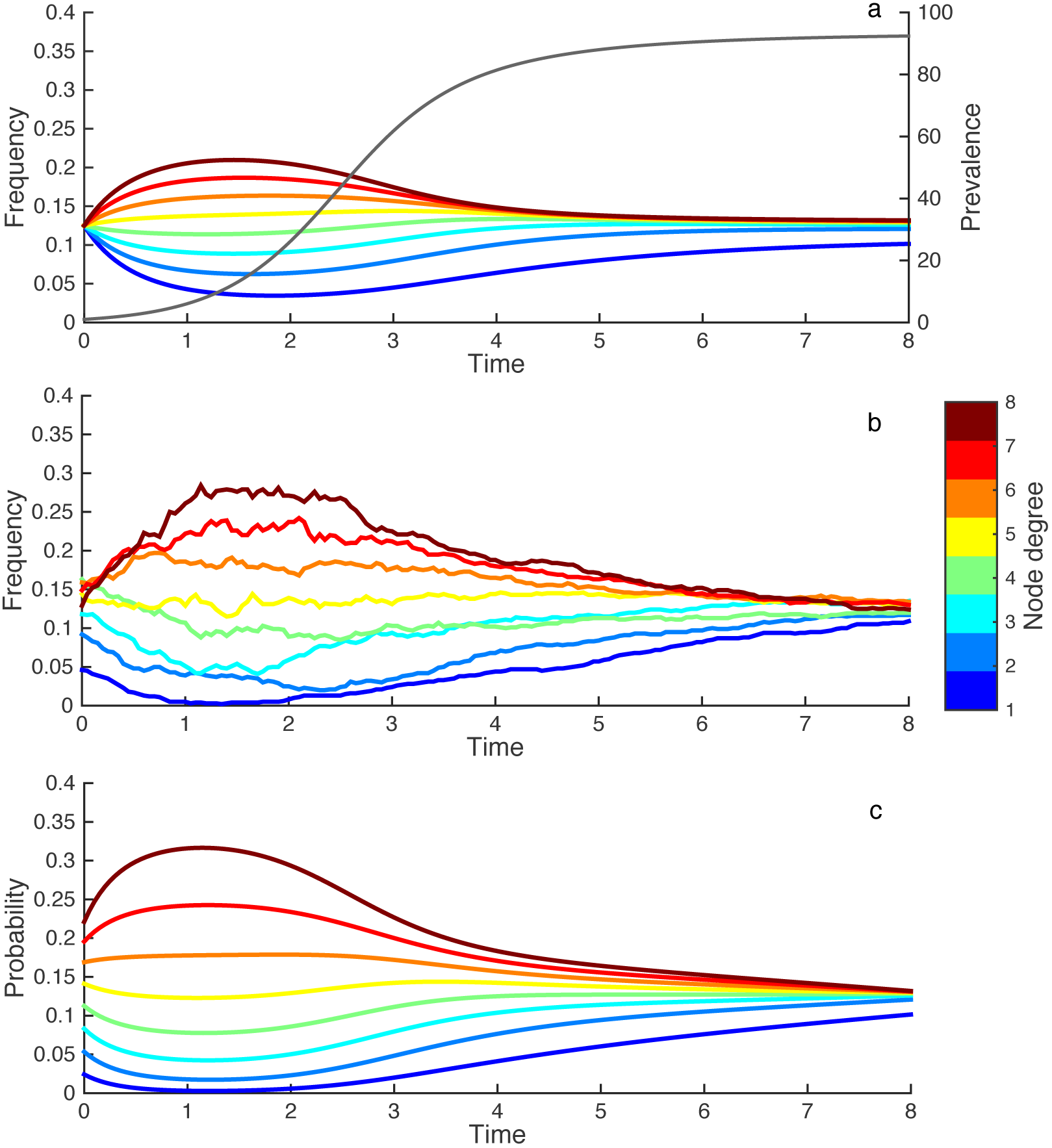
The degree distribution of infected nodes and ancestral lineages over the time course of an epidemic. **(a)** Degree distribution for the entire infected population. At time *t* = 0 we start with a uniform degree distribution for the initial infected host that mirrors the uniform degree distribution of the underlying network. The overlaid grey line represents prevalence over time. **(b)** The ancestral degree distribution for a single lineage traced backwards through time from 1000 stochastic simulations **(c)** Theoretical expectation for the ancestral degree of a single lineage given by the master equations derived from the pairwise model.

### Simulation methods

For each individual-based stochastic (IBS) simulation, we first generated a random network with the desired statistical properties using the configuration model [30]. If the network was not completely connected with all nodes connected to all others by at least one path in the network, it was discarded and a new one generated. To seed the epidemic, a single node was then randomly chosen to become infected at time *t* = 0. The simulations then preceded forwards in time using an event-driven approach similar to the Gillespie stochastic simulation algorithm [47]. Infected hosts were allowed to either transmit to their susceptible neighbors or recover from infection. At transmission events, the parent and child pathogen lineage were recorded so that the ancestry of each lineage could be traced in order to recover the true phylogeny of the pathogen population. At recovery events, infected individuals were sampled with probability *ρ* and subsequently included in the phylogeny. At the final time *t* = *T*, all surviving infections were also sampled with probability *ρ* and included in the phylogeny. Unless otherwise stated, infected individuals were sampled serially though time upon removal (i.e. recovery).

Phylogenies were simulated under the pairwise coalescent model backwards in time. At time *t* = *T*, sampled individuals were added to a set of lineages that were then traced back through time. The degree of sampled lineages was drawn randomly from the degree distribution of the infected population at the time of sampling according to the pairwise epidemic model. The state of each lineage was then updated incrementally using small time steps receding into the past. At each discrete time step, lineages could either transition to a different degree node or coalesce with another lineage with probabilities proportional the rate of transitions and coalescent events given Eq (16) above and Eq (12) in the main text. Simulations were run until the final two ancestral lineages coalesced.

### When the pairwise approximation fails

From the general theory of dynamical processes on networks, we expect pair approximations to work well when dynamical correlations arise locally at the level of pairs or other lower-order motifs like triples, but may break down when there is significant higher-order network structure, such as when the network is modular or broken up into different communities [34, 48]. Given that the coalescent process can also be viewed as a dynamical process on a network (albeit backwards in time), we expect that the pair approximations underlying the pairwise coalescent will also break down in the presence of higher-order network structure. To explore how higher-order structure affects the accuracy of the pairwise models, we used the well-known Watts-Strogatz model [49] to generate networks with varying levels of higher-order structure.

To simulate random graphs under the Watt-Strogatz model we start with an initially perfectly ordered ring network where each node is connected to its 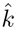 nearest neighbors and then randomly rewire a fraction of edges *f*. A low *f* therefore preserves the original community structure present in the ring whereas a high *f* randomizes the network in a way that destroys higher-order structure (SI Fig 2). Our variant of this algorithm uses degree-preserving rewiring so that we can study the effects of community structure without introducing additional contact heterogeneity.

**SI Figure 2.**
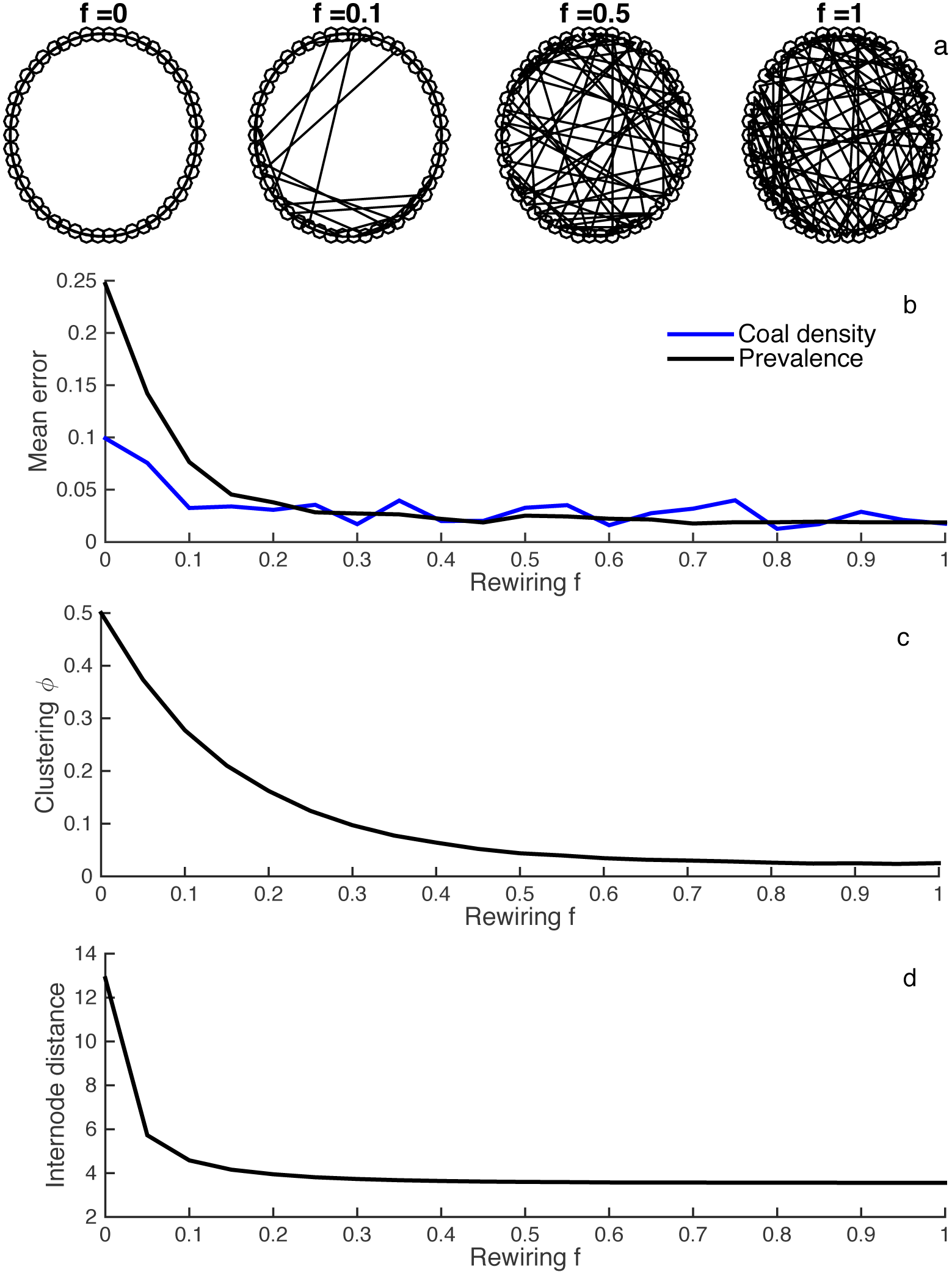
Accuracy of pairwise approximation on Watts-Strogatz networks with varying levels of higher-order community structure. **(a)** Watts-Strogatz networks with different edge rewiring fractions *f*. For ease of viewing, *N* = 50 here. **(b)** Time-integrated mean error in the coalescent density (blue) and prevalence (black) given by the pairwise models when compared against stochastic simulations on networks randomly generated with different *f* values. (**c**) The mean clustering coefficient *ϕ* of simulated networks for each *f* value. (**d**) The mean internode distance for the same networks as in b and c. For all simulations 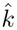 = 4 and *N* = 250.

To measure the error in the theoretical expectations provided by the pairwise models when compared against individual-based stochastic (IBS) simulations on Watts-Strogatz networks, we approximate the time-integrated mean error *Ē* in prevalence and coalescent distributions by averaging over all *T* time points on the discretized interval, such that

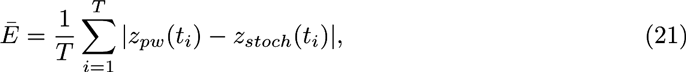

where *z_pw_* is the value given by the pairwise model and *z_stoch_* is the mean value given by the stochastic simulations.

The time-integrated mean error in the theoretical expectations provided by the pairwise models for both prevalence and the distribution of coalescent events is shown in SI Fig 2. The error arising from the pair approximation is only large when the rewiring fraction is very low (*f* <= 0.10) and there is substantial community structure in the networks. Moreover, the pairwise coalescent model appears to break down at the same point as the pairwise epidemic model, which is not surprising given that the coalescent model depends on the accuracy of the epidemic dynamics predicted by the pairwise epidemic model. While networks with *f* <= 0.10 have high clustering coefficients, this does not appear to be the ultimate downfall of the pairwise models because there is already substantial clustering with *f* > 0.10 where the pairwise models still perform well (SI Fig 2). Rather, where the pairwise models breaks down at *f* <= 0.10 is also the point at which we see a large spike in the mean internode distance, the minimum distance between two nodes in a network (SI Fig 2).

Thus, the pairwise models perform well as long as the networks are sufficiently “small” as quantified by mean internode distances, which will rise sharply once the network is broken up into different communities. This echoes an earlier observation made by [48], who showed low-dimensional models that ignore higher-order community structure can provide surprisingly accurate approximations to dynamics on a variety of complex networks as long as networks are sufficiently small-world.

### Statistical performance of estimators

The statistical performance of the pairwise coalescent model in estimating network properties from phylogenies was extensively tested on simulated phylogenies. To check for potential biases in our estimates of network connectivity, clustering, contact heterogeneity and assortativity, we simulated 100 additional phylogenies under a fixed value of each parameter using forward-time IBS simulations. We then obtained a maximum likelihood estimate (MLE) of the corresponding parameter from each tree using a nonlinear numerical optimization routine. All other epidemiological parameters were fixed at their true values. The MLEs appear centered around the true parameter values with little to no detectable bias (SI Fig 3).

Next, we simulated trees under a wider range of parameter values for each network property to check how well our estimator performs under different model parameterizations. Overall, parameter estimates appear well-calibrated with a high correlation between the true and MLE values (SI Fig 3). While the coverage of our confidence intervals falls below the desired 95% level, we believe the coverage achieved is very reasonable given that the pairwise models ignore stochastic variation in both network topology and epidemic dynamics, which can cause tree structure to diverge considerably from what is theoretically expected under the pairwise models.

**SI Figure 3.**
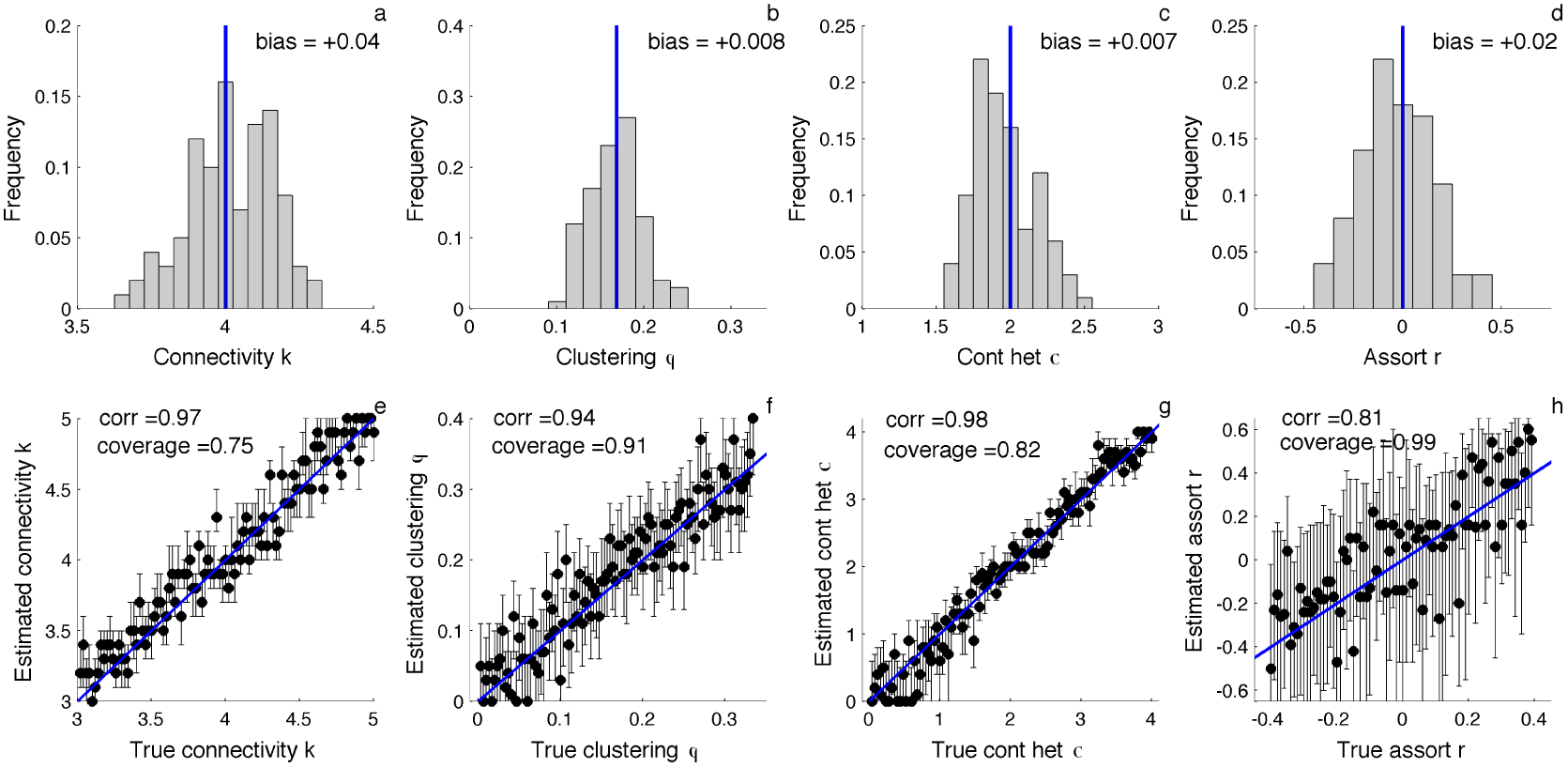
Likelihood-based estimates of network parameters and their statistical performance. **(a-d)** Distribution of maximum likelihood point estimates of the overall connectivity 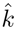 clustering coefficient *ϕ*, contact heterogeneity *σ_k_* and assortatvity coefficient *r* for 100 simulated phylogenies. (**e-h**) Maximum likelihood estimates (dots) and 95% confidence intervals (lines) for each network property under different model parameterizations. Blue lines indicate the true parameter value used in each simulation. Correlation (corr) refers to the Pearson correlation between the true and estimated parameter values. Coverage refers to the fraction of simulations in which the true parameter falls within the estimated 95% confidence intervals. All simulations were performed with a fraction *ρ* = 0.5 of infected individuals sampled upon removal serially over time.

### Phylogenetic analysis of Swiss HIV-1 sequence data

The HIV-1 subtype B epidemic in Switzerland (hereafter CH) is strongly integrated into the general European subtype B epidemic, especially among MSM [42, 50]. We therefore first tried to identify sub-epidemics primarily occurring on local contact networks within CH rather than abroad. We combined 4441 subtype B *pol* sequences taken from MSM patients enrolled in the Swiss HIV Cohort Study (SHCS) with a large background dataset of 4550 subtype B sequences from the Los Alamos National Laboratory (LANL) HIV database. After removing all non-subtype B and recombinant sequences, the SHCS and LANL sequences were then aligned together against the HBX2 subtype B reference strain. After alignment, a total of 51 codon positions associated with known drug resistance mutations were also stripped from the alignment. A maximum likelihood (ML) phylogeny of the combined LANL + SHCS alignment was then reconstructed in FastTree [51] assuming a GTR model of molecular evolution with gamma distributed rate heterogeneity.

To identify sub-epidemics occurring predominantly within CH, we first reconstructed the ancestral location of all internal nodes using maximum parsimony. Introductions into CH were assumed to occur whenever a node inferred to be in CH had a parent node outside of CH. Sub-epidemics were then defined to include all lineages sampled in CH that descended from an introduction event into CH without passing through a node reconstructed to be outside of CH. This preliminary analysis revealed that the Swiss epidemic is composed of many sub-epidemics likely originating from independent introductions into CH. Most of these sub-epidemics are composed of only a few sampled individuals and can be categorized as occurring predominantly in either the French or German speaking region of CH (SI Fig 4). To minimize the effects of geographic structure within CH on our phylodynamic analysis, we chose to focus on a large cluster which included 200 sampled individuals who predominantly lived or sought treatment in the Zürich area.

**SI Figure 4.**
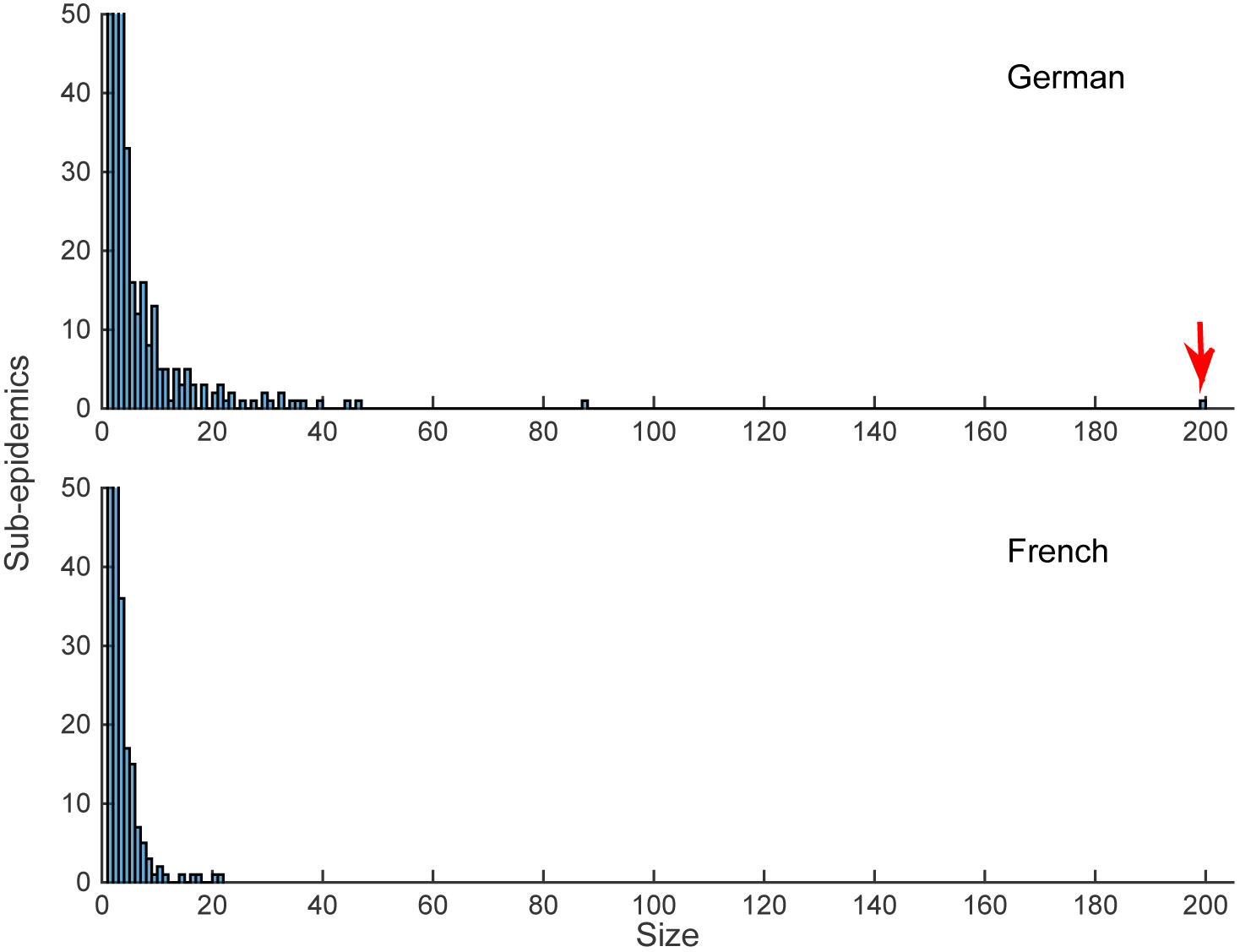
Size distribution of HIV sub-epidemics in Switzerland. Sub-epidemics were categorized as occurring in either the German or French speaking regions of Switzerland. Size refers to the number of sampled individuals included in each sub-epidemic. The red arrow marks the large sup-epidemic we chose to analyze in detail.

From these 200 sequences, we reconstructed a new time-calibrated phylogeny in BEAST 2 assuming a strict molecular clock, a GTR substitution model and a Bayesian Skyline prior on effective population size through time [29]. The maximum clade credibility tree from this initial BEAST analysis was then fixed for our phylodynamic analysis using the pairwise coalescent model. To this tree, we fit a SIR-type pairwise epidemic model where the degree distribution was modeled as a discretized gamma distribution. In general, we used fairly informative priors on the epidemiological parameters in the model but relatively uninformative priors on the network parameters (SI Table 1). This model was implemented in BEAST 2 as an add-on package called PairTree, freely available at https://github.com/davidrasm/PairTree. Posterior distributions for all model parameters were inferred using BEAST’s built-in MCMC sampling algorithm.

**Table 1.**
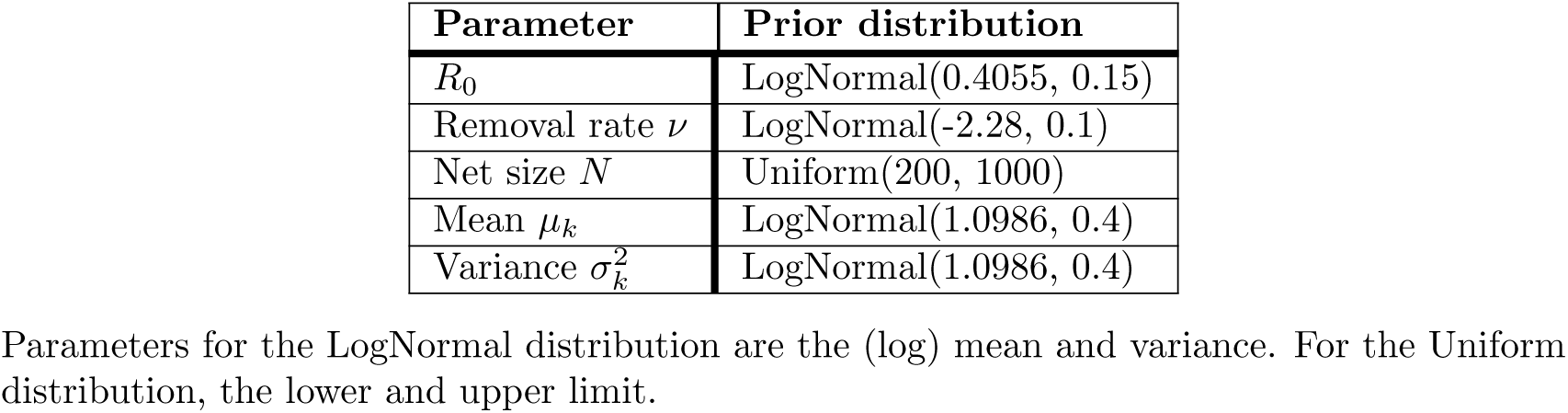
Priors on the parameters used in the Swiss HIV analysis.

## References

1. Garnett GP, Hughes JP, Anderson RM, Stoner BP, Aral SO, Whittington WL, et al. Sexual mixing patterns of patients attending sexually transmitted diseases clinics. Sexually Transmitted Diseases. 1996;23(3):248–257.

2. Helleringer S, Kohler HP. Sexual network structure and the spread of HIV in Africa: evidence from Likoma Island, Malawi. AIDS. 2007;21(17):2323–2332.

3. Liljeros F, Edling CR, Amaral LAN, Stanley HE, Åberg Y. The web of human sexual contacts. Nature. 2001;411(6840):907–908.

4. Rocha LE, Liljeros F, Holme P. Simulated epidemics in an empirical spatiotemporal network of 50,185 sexual contacts. PLoS Comput Biol. 2011;7(3):e1001109.

5. Newman ME. Spread of epidemic disease on networks. Physical Review E. 2002;66(1):016128.

6. Pastor-Satorras R, Vespignani A. Immunization of complex networks. Physical Review E. 2002;65(3):036104.

7. Keeling MJ, Eames KT. Networks and epidemic models. Journal of the Royal Society Interface. 2005;2(4):295–307.

8. Meyers LA, Pourbohloul B, Newman ME, Skowronski DM, Brunham RC. Network theory and SARS: predicting outbreak diversity. Journal of Theoretical Biology. 2005;232(1):71–81.

9. Bansal S, Grenfell BT, Meyers LA. When individual behaviour matters: homogeneous and network models in epidemiology. Journal of the Royal Society Interface. 2007;4(16):879–891.

10. Liljeros F, Edling CR, Amaral LAN. Sexual networks: implications for the transmission of sexually transmitted infections. Microbes and Infection. 2003;5(2):189–196.

11. Eames K, Bansal S, Frost S, Riley S. Six challenges in measuring contact networks for use in modelling. Epidemics. 2015;10:72–77.

12. Yerly S, Vora S, Rizzardi P, Chave JP, Vernazza P, Flepp M, et al. Acute HIV infection: impact on the spread of HIV and transmission of drug resistance. AIDS. 2001;15(17):2287–2292.

13. Ragonnet-Cronin M, Ofner-Agostini M, Merks H, Pilon R, Rekart M, Archibald CP, et al. Longitudinal phylogenetic surveillance identifies distinct patterns of cluster dynamics. Journal of Acquired Immune Deficiency Syndromes. 2010;55(1):102–108.

14. Brown AJL, Lycett SJ, Weinert L, Hughes GJ, Fearnhill E, Dunn DT. Transmission network parameters estimated from HIV sequences for a nationwide epidemic. Journal of Infectious Diseases. 2011;204(9):1463–1469.

15. Leitner T, Escanilla D, Franzen C, Uhlen M, Albert J. Accurate reconstruction of a known HIV-1 transmission history by phylogenetic tree analysis. Proceedings of the National Academy of Sciences. 1996;93(20):10864–10869.

16. Cottam EM, Thébaud G, Wadsworth J, Gloster J, Mansley L, Paton DJ, et al. Integrating genetic and epidemiological data to determine transmission pathways of foot-and-mouth disease virus. Proceedings of the Royal Society of London B: Biological Sciences. 2008;275(1637):887–895.

17. Jombart T, Eggo R, Dodd P, Balloux F. Reconstructing disease outbreaks from genetic data: a graph approach. Heredity. 2011;106(2):383–390.

18. Ypma R, Bataille A, Stegeman A, Koch G, Wallinga J, Van Ballegooijen W. Unravelling transmission trees of infectious diseases by combining genetic and epidemiological data. Proceedings of the Royal Society of London B: Biological Sciences. 2012;279(1728):444–450.

19. Didelot X, Gardy J, Colijn C. Bayesian inference of infectious disease transmission from whole-genome sequence data. Molecular Biology and Evolution. 2014;31(7):1869–1879.

20. Hall M, Woolhouse M, Rambaut A. Epidemic Reconstruction in a Phylogenetics Framework: Transmission Trees as Partitions of the Node Set. PLoS Comput Biol. 2015;11(12):e1004613.

21. O’Dea EB, Wilke CO. Contact heterogeneity and phylodynamics: how contact networks shape parasite evolutionary trees. Interdisciplinary Perspectives on Infectious Diseases. 2010;2011.

22. Leventhal GE, Kouyos R, Stadler T, Von Wyl V, Yerly S, Böni J, et al. Inferring epidemic contact structure from phylogenetic trees. PLoS Comput Biol. 2012;8(3):e1002413–e1002413.

23. Robinson K, Fyson N, Cohen T, Fraser C, Colijn C. How the dynamics and structure of sexual contact networks shape pathogen phylogenies. PLoS Comput Biol. 2013;9(6):e1003105.

24. Volz EM, Koopman JS, Ward MJ, Brown AL, Frost SD. Simple epidemiological dynamics explain phylogenetic clustering of HIV from patients with recent infection. PLoS Comput Biol. 2012;8(6):e1002552–e1002552.

25. Newman M. Networks: An Introduction. Oxford University Press; 2010.

26. Keeling MJ. The effects of local spatial structure on epidemiological invasions. Proceedings of the Royal Society of London B: Biological Sciences. 1999;266(1421):859–867.

27. Rand D. Correlation equations and pair approximations for spatial ecologies. Advanced Ecological Theory: Principles and Applications. 1999;100.

28. Eames KT, Keeling MJ. Modeling dynamic and network heterogeneities in the spread of sexually transmitted diseases. Proceedings of the National Academy of Sciences. 2002;99(20):13330–13335.

29. Bouckaert R, Heled J, Kühnert D, Vaughan T, Wu CH, Xie D, et al. BEAST 2: a software platform for Bayesian evolutionary analysis. PLoS Comput Biol. 2014;10(4):e1003537.

30. Molloy M, Reed BA. A critical point for random graphs with a given degree sequence. Random Structures and Algorithms. 1995;6(2/3):161–180.

31. Miller JC. Percolation and epidemics in random clustered networks. Physical Review E. 2009;80(2):020901.

32. Newman ME. Random graphs with clustering. Physical Review Letters. 2009;103(5):058701.

33. Newman ME. Mixing patterns in networks. Physical Review E. 2003;67(2):026126.

34. Taylor M, Simon PL, Green DM, House T, Kiss IZ. From Markovian to pairwise epidemic models and the performance of moment closure approximations. Journal of Mathematical Biology. 2012;64(6):1021–1042.

35. Volz EM, Pond SLK, Ward MJ, Brown AJL, Frost SD. Phylodynamics of infectious disease epidemics. Genetics. 2009;183(4):1421–1430.

36. Frost SD, Volz EM. Viral phylodynamics and the search for an ‘effective number of infections’. Philosophical Transactions of the Royal Society B: Biological Sciences. 2010;365(1548):1879–1890.

37. Koelle K, Rasmussen DA. Rates of coalescence for common epidemiological models at equilibrium. Journal of The Royal Society Interface. 2012;9(70):997–1007.

38. Volz EM. Complex population dynamics and the coalescent under neutrality. Genetics. 2012;190(1):187–201.

39. House T, Keeling MJ. Insights from unifying modern approximations to infections on networks. Journal of The Royal Society Interface. 2011;8(54):67–73.

40. Frost SD, Volz EM. Modelling tree shape and structure in viral phylodynamics. Philosophical Transactions of the Royal Society of London B: Biological Sciences. 2013;368(1614):20120208.

41. Ledergerber B, Egger M, Opravil M, Telenti A, Hirschel B, Battegay M, et al. Clinical progression and virological failure on highly active antiretroviral therapy in HIV-1 patients: a prospective cohort study. The Lancet. 1999;353(9156):863–868.

42. Kouyos RD, Von Wyl V, Yerly S, Böni J, Taffé P, Shah C, et al. Molecular epidemiology reveals long-term changes in HIV type 1 subtype B transmission in Switzerland. Journal of Infectious Diseases. 2010;201(10):1488–1497.

43. Marzel A, Shilaih M, Yang WL, Böni J, Yerly S, Klimkait T, et al. HIV-1 Transmission During Recent Infection and During Treatment Interruptions as Major Drivers of New Infections in the Swiss HIV Cohort Study. Clinical Infectious Diseases. 2016;62(1):115–122.

44. Swiss Federal Office of Public Health. HIV and AIDS in der Schweiz. 2011;.

45. Bauch C, Rand D. A moment closure model for sexually transmitted disease transmission through a concurrent partnership network. Proceedings of the Royal Society of London B: Biological Sciences. 2000;267(1456):2019–2027.

46. Eames KT, Keeling MJ. Monogamous networks and the spread of sexually transmitted diseases. Mathematical Biosciences. 2004;189(2):115–130.

47. Gillespie DT. Stochastic simulation of chemical kinetics. Annu Rev Phys Chem. 2007;58:35–55.

48. Melnik S, Hackett A, Porter MA, Mucha PJ, Gleeson JP. The unreasonable effectiveness of tree-based theory for networks with clustering. Physical Review E. 2011;83(3):036112.

49. Watts DJ, Strogatz SH. Collective dynamics of ‘small-world’ networks. Nature. 1998;393(6684):440–442.

50. Paraskevis D, Pybus O, Magiorkinis G, Hatzakis A, Wensing AM, Vijver DA, et al. Tracing the HIV-1 subtype B mobility in Europe: a phylogeographic approach. Retrovirology. 2009;6(1):1.

51. Price MN, Dehal PS, Arkin AP. FastTree 2–approximately maximum-likelihood trees for large alignments. PloS One. 2010;5(3):e9490.

